# A regulatory RNA is associated to invasive meningococcal disease in Europe

**DOI:** 10.1101/583476

**Authors:** Jens Karlsson, Hannes Eichner, Susanne Jacobsson, Edmund Loh

## Abstract

The strictly human pathogen *Neisseria meningitidis* is a commensal bacterium but can occasionally turn lethal causing septicaemia and meningitis. The mechanisms of how the meningococcus shifts to invasive infection remain poorly understood. Here we demonstrate that an eight base-pair tandem repeat deletion in the 5’-untranslated region of the polysaccharide capsular biosynthesis operon results in a hypercapsulation phenotype in clinical isolates. The increased capsule production significantly improves the bacterium survival in human serum while impairing its ability to adhere and colonise human pharyngeal cells. Among 4501 reported meningococcal cases in Europe from 2010-2018, the loss of an eight base-pair tandem repeat is three times more prevalent in invasive isolates (16.3%) compared to carrier isolates (5.1%).Combined results indicate that polymorphisms in this regulatory RNA contributes to meningococcal virulence.

**Importance:** In this study we report a regulatory RNA to be directly involved in clinical manifestation of meningococcal disease. Using readily accessible WGS of meningococcus, we have now demonstrated that regulatory RNAs directly contribute to the progression of invasive meningococcal infection. We believe this novel combination of molecular and comparative regulatory RNA study could be used for the identification of additional RNAs involved in not only meningococcus but also pave the way for similar studies in other important bacterial pathogens. The identification of specific regulatory RNAs will no doubt facilitate clinicians, microbiologists, and public health practitioners to adjust their diagnostic techniques and treatments to best fit the condition of the patients.

## Introduction

*Neisseria meningitidis* is a Gram-negative diplococcus and an obligate human commensal bacterium, which resides exclusively on the epithelium of the nasopharynx. Carriage is age-dependent and rises from 4.5% in infants (0-1 years) to 23.7% in adolescents (15-19 years) and down to 7.8% in middle aged adults (50 years) (1). By mechanisms not fully understood, the harmless colonisation can rapidly turn into an invasive infection leading to lethal septicaemia and meningitis. The incidence of invasive meningococcal disease is highly dependent on geographical region and season, and ranges from 0.1/100,000/population/year to almost 100/100,000/population/year (2). People with increased risk for invasive meningococcal infections are infants, adolescents and immunocompromised individuals (3-5).

Currently, there are twelve known serogroups of *N. meningitidis* identified based on the composition of their capsular polysaccharide (A, B, C, E – H, I – K, L – W, X, Y, Z) with six of these responsible for outbreaks; A, B, C, W, X, and Y (6). The *capsular synthesis* (*cps*) locus has been characterised for a wide range of serogroups and shows gene order synteny (6). In addition to serogroup, meningococci can be grouped into clonal complexes (cc) and sequence types (ST) by multilocus sequence typing (MLST) based on polymorphisms in housekeeping genes (7). In this study, we investigated meningococcal isolates from two Swedish adolescents who contracted meningococci after attending a ski trip to the French Alps in March 2017. During the trip one adolescent contracted invasive meningococceamia and recovered. Two months later after returning home to Sweden, another adolescent succumbed to invasive meningococceamia and a third adolescent was identified as an asymptomatic carrier. This study aims to elucidate and compare virulence mechanisms underlying the cause for the fatal meningococcal infection especially in the regulation of its polysaccharide capsule production. Using findings from the Swedish isolates, the subsequent work encompasses the investigation of correlations in the manifestation of diseases between 4501 meningococcal invasive- and carrier isolates from Europe.

## Results

We whole-genome sequenced (WGS) both the Swedish meningococcal isolates used in this study and deposited them in the PubMLST database under following IDs; 17-264: 53777 (invasive isolate) and 17-271: 53778 (carrier isolate). Both isolates were identified to belong to serogroup C, cc32 and with identical typing genes as the French isolate (Table 1). Bacterial growth, micro-, and macroscopical examinations revealed no significant difference between the isolates (Fig S1 and S2). Loci similarity comparison from the WGS encompassing 2219 loci revealed 98.3% (2181) loci conservation (Table EV2). The 1.7% (38) different loci between the isolates comprise of incomplete *IS1655* transposase insertions in the 17-271 isolate, together with genes coding for outer membrane proteins, efflux pumps, and metabolic proteins (Table 3). Further analyses revealed that many hits of the variable loci where due to single nucleotide polymorphisms (SNPs) that were either true SNPs or sequencing errors. All true SNPs identified were found to generate the same gene products in both isolates. In addition, other hits were also later identified as sequencing error due to long Poly-G or Poly-C regions present over the locus.

**Table 1.**
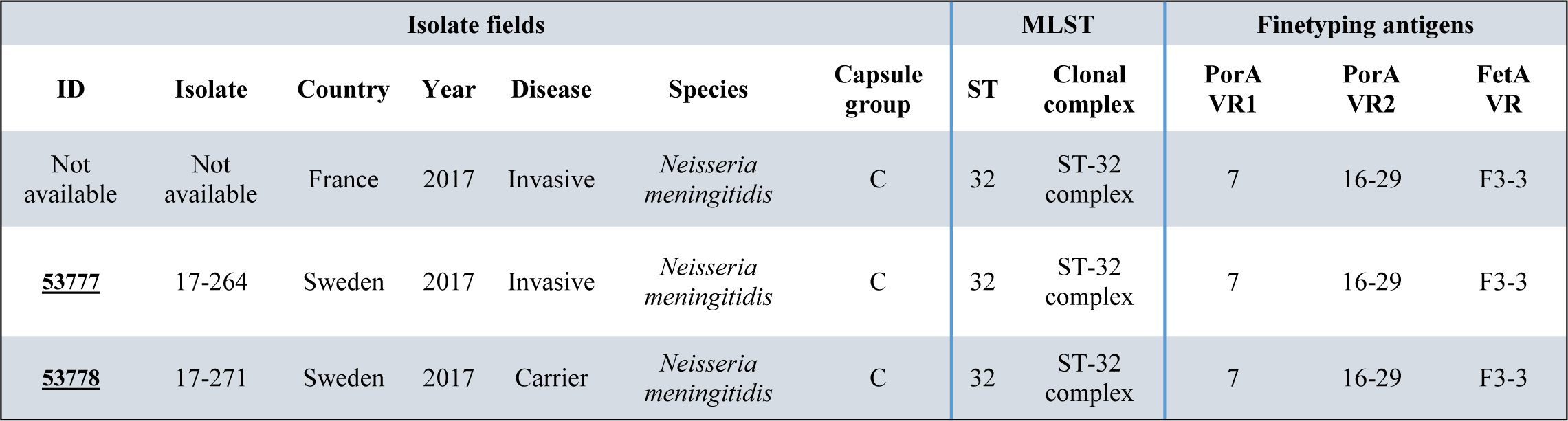
Typing output of 17-264 and 17-271 isolates recovered from PubMLST as well as the typing of the isolate responsible for the infection in the French Alps.

**Table 2.** Summary of loci comparison between 17-264 and 17-271 isolates. Missing-, and incomplete loci in both isolates were excluded from calculations.

| Matching loci | Missing loci | Different loci | Incomplete loci | Frequency matching (%) |
| --- | --- | --- | --- | --- |
| 2181 | 763 | 38 | 60 | 98,3 |

**Table 3.** List of the 38 different loci in the WGS comparison between 17-264 and 17-271.

| Locus | Product | Cause/Effect |
| --- | --- | --- |
| NEIS0210 | *pilE* | Antigenic variation |
| NEIS0568 | *pglE* | 7bp tandem repeats (17-264: 11 and 17-271: 13), different frame but both off phase |
| NEIS2986 | hypothetical protein | Different, pseudogene |
| igr_up_NEIS0055 | 5'-UTR-*cssA* | 8bp tandem repeat missing in 17-264 |
| NEIS1310 | *modA12* | 4bp tandem repeats (17-264: 14 and 17-271: 23), same frame and both off phase |
| NEIS0033 | *pilC2* | Poly-G region sequencing error.<br>WB shows no difference |
| NEIS0054 | *cssA* | SNP |
| NEIS0085 | protein export protein | End of contig* in 17-271 sequencing but otherwise identical |
| NEIS0213 | *pglA* | SNP/undecided nucleotide in sequencing |
| NEIS0297 | insertion element *IS1655* transposase | Incomplete transposase element in 17-271 |
| NEIS0377 | putative integral membrane protein | SNP/undecided nucleotide in sequencing |
| NEIS0380 | *pglL* | End of contig* in 17-264 sequencing but otherwise identical |
| NEIS0402 | *pglF* | End of contig* in 17-264 sequencing but otherwise identical |
| NEIS0493 | hypothetical protein | 5' sequencing error, undecided nucleotides |
| NEIS0648 | *ksgA* | SNP/undecided nucleotide in sequencing |
| NEIS0710 | *pnp* | SNP |
| NEIS0715 | cysteine synthase | SNP |
| NEIS0904 | transposase for *IS1655* | Incomplete transposase element in 17-271 |
| NEIS0976 | transposase for *IS1655* | Incomplete transposase element in 17-271 |
| NEIS1008 | transposase for *IS1655* | Incomplete transposase element in 17-271 |
| NEIS1055 | transposase for *IS1655* | Incomplete transposase element in 17-271 |
| NEIS1085 | *mpl* | SNP |
| NEIS1156 | hypothetical protein | SNP, Poly-G region sequencing error |
| NEIS1288 | putative aldehyde dehydrogenase | SNP and end of contig* |
| NEIS1381 | transposase for *IS1655* | Incomplete transposase element in 17-271 |
| NEIS1418 | putative membrane peptidase | Sequencing error |
| NEIS1852 | *farA* | Sequencing error and end of contig* |
| NEIS1853 | *farB* | End of contig* |
| NEIS1884 | transposase for *IS1655* | Incomplete transposase element in 17-271 |
| NEIS1902 | *lgtA* | Poly-G region sequencing error and end of contig* |
| NEIS1926 | putative inner membrane protein | End of contig* |
| NEIS1943 | *nalP* | SNP, Poly-C region sequencing error in 17-271 |
| NEIS1993 | insertion element *IS1655* transposase | Incomplete transposase element in 17-271 |
| NEIS2099 | putative immunity protein | Sequencing error and end of contig* |
| NEIS2124 | lipoprotein | End of contig* |
| NEIS2125 | insertion element *IS1655* transposase | Incomplete transposase element in 17-271 |
| NEIS2155 | *lgtD* | Poly-G region sequencing error and end of contig* |
| igr_up_NEIS1364 | *porA* promoter | SNP in 17-271.<br>WB shows no difference |

\*End of contig refers to sequencing contig is incomplete in covering the loci in one WGS sample and therefore are in the list as different. Unless stated otherwise the sequencing is identical up to that point

The final five hits confirmed as different are NEIS0210 (*pilE*), IGR^up^NEIS0055 (5’-UTR-*cssA*), NEIS0568 (*pglE*), NEIS1310 (*modA12*), and NEIS2986 (hypothetical protein) (Highlighted in Table EV3). The *pilE* gene encodes for the major subunit of Type-IV pilus (T4P) and is different due to antigenic variation, usually observed even within the same clonal population (8). The 5’-untranslated region (UTR) of the capsular biosynthesis operon *cssA* of the invasive isolate 17-264 lacks an eight base pair (8bp) tandem repeat (5’-TATACTTA-3’). The *pglE* gene is involved in the glycosylation of T4P and is known for its ON/OFF phase-variation depending on heptarepeats (5’-CAAACAA-3’) (9). The two isolates have different transcriptional *pglE* frames due to different amount of heptarepeats but both result in the same OFF phase. Similar to the regulation to *pglE*, the *modA12* gene encoding a methyltransferase of DNA is regulated by tetrarepeats, 5’-AGCC-3’ (10) and although both isolates have different amounts of repeats, they are both in the same OFF phase. NEIS2986 is a hypothetical protein of 90 amino acid, with no elucidated function and no recognised protein motifs. The predicted protein sequence does not contain start codon, which indicate that this is a pseudogene. Overall, among the 38 loci identified as different, the only *bona fide* hit is the 5’-UTR-*cssA* between the two isolates.

Through immunoblot analysis, we reveal that the invasive isolate 17-264 has a higher expression of CssA (capsular synthesis protein), PilE (predominant minor pilin in the T4P involved in adhesion), and Opa,(opacity proteins involved in adhesion) (Fig 1A). Expression of other virulence factors such as fHbp, (responsible for sequestering immune factor H), Hfq (RNA-chaperone important for sRNA-mediated gene regulation), PorA (voltage-gated, cation channel) RmpM (periplasmic protein interacting and stabilising PorA and PorB), and other pilin proteins (pilus and pilus-associated proteins that form the T4P) remain same for both isolates and were used as controls. The capsular polysaccharide *cssA* coding region was also initially annotated as different between the isolates (Table EV3), but further examination reveals the presence of a single base polymorphism but coding for the same amino acid and therefore does not explain the higher expression of CssA in invasive isolate 17-264 compared to carrier isolate 17-271. These observations prompted us to further investigate into the regulatory region of the capsular polysaccharide operon.

**Fig. 1.**
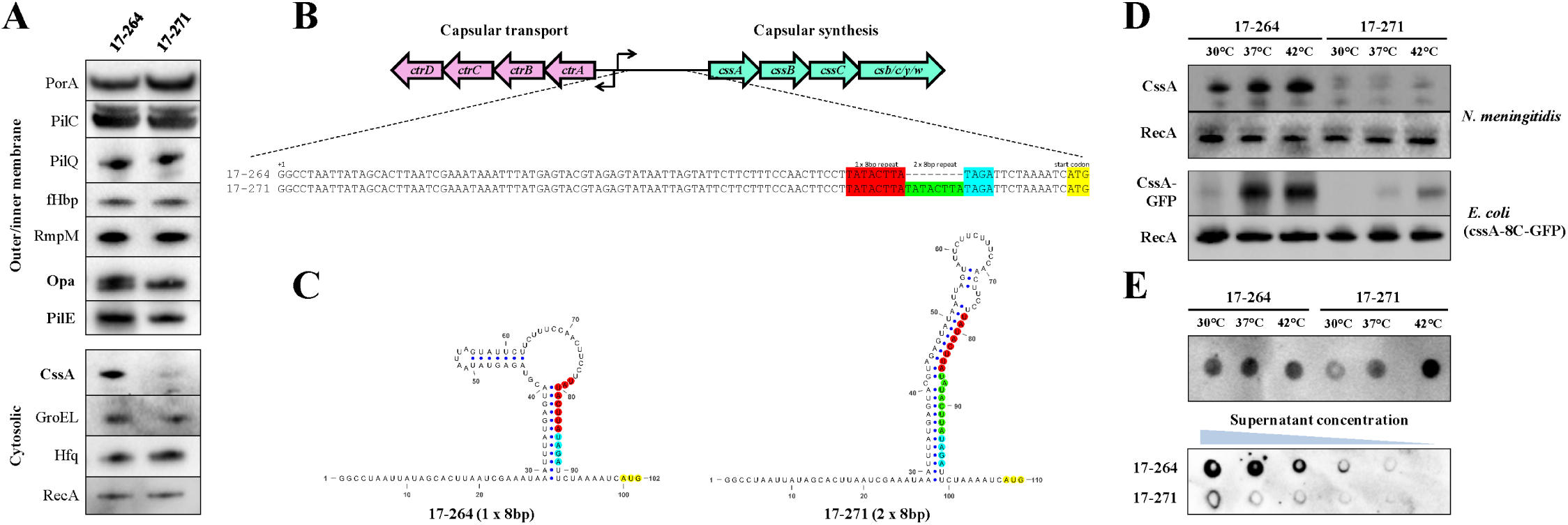
5’-UTR-cssA mapping, virulence factor characterisation, and temperature-mediated expression of CssA and capsule. A) Immunoblot panel display the expression level of a variety of virulence related-and house-keeping proteins. Highlighted in bold are the virulence proteins upregulated in the invasive isolate 17-264; CssA, Opa and PilE. B) Illustration showing the ctr and cps loci with focus on the intergenic region. The 5’-UTR-cssA sequences from the two clinical isolates were aligned. An 8bp deletion was observed in the invasive isolate.C) Secondary structure prediction of the 5’-UTR-cssA RNA using VARNA14 illustrating the thermodynamic stable stem-loop structure of the carrier isolate 17-271 (ΔG: −15.60 kcal/mol) and a smaller and less stable stem-loop of the invasive isolate 17-262 (ΔG: −11.30 kcal/mol). The tandem repeat, RBS and start codon are marked in red/green, cyan and yellow, respectively. Small blue dots designate base pairing. D) Immunoblots of temperature mediated expression shows an increasing and constitutive high expression of CssA in the invasive isolate 17-264 compared to carrier isolate 17-271 (top). The 8bp tandem repeats function ectopically, controlling EGFP expression similar to their respective meningoccal background. (lower). E) Dot blots of capsular production on the surface of the bacteria show a constitutively high capsule production in the invasive isolate 17-264 compared to carrier isolate 17-271 (top). Serial dilution of supernatant from liquid culture grown at 37°C shows that the invasive isolate 17-264 sheds more capsule into the medium compared to the carrier isolate 17-271 (lower). Data information: All blots are representative of at least three biological replicates.

The WGS data reveal the loss of an 8bp tandem repeat in the 5’-UTR-*cssA* region of the invasive isolate 17-264 compared to the carrier isolate 17-271 (Fig 1B). Secondary structure predictions of the 5’-UTR-*cssA* mRNA confirms a possible stem-loop formation in the carrier isolate 17-271 (*ΔG*: −15·60 kcal/mol), while the invasive isolate 17-264 possess a thermodynamically less stable (*ΔG*: −11·30 kcal/mol) secondary structure (Fig 1C). Previously, we identified that the 5’-UTR-*cssA*, functions as an RNA-.thermosensor, controlling the expression of CssA in a temperature dependent manner (11). We therefore investigated temperature dependent expression of CssA in these two clinical isolates. Results show a temperature-mediated upregulation of CssA in the 17-264 whereas the level of CssA in the 17-271 remained low (Fig 1D - top). Since a *bona fide* RNA-thermosensor should be able to function independently of the native bacterial host factors, an ectopic bacterial host (*Escherichia coli*) was selected to further investigate this thermal regulation. The respective 5’-UTR-*cssA* from each isolate was introduced upstream of a green fluorescent reporter EGFP in the pEGFP-N2 vector and transformed into *E. coli*. Immunoblot results from *E. coli* were consistent with their respective meningococcal background (Fig 1D – lower).

An elevated expression of the CssA protein is however not definite evidence that more surface capsule is produced. To address this, “dot blots” assays were performed, as it has previously been used to semi-quantitatively measure meningococcal capsule production (12-14). Dot blots using serogroup C specific anti-sera show that surface polysaccharide capsule is produced in high amounts regardless of cells, Detroit 562 were performed. Results reveal that the less capsulated carrier 17-271 isolate has a significant, ten times more effective adherence to the cells compared to the hypercapsulated invasive 17-264 isolate (Fig 2B).

**Fig. 2.**
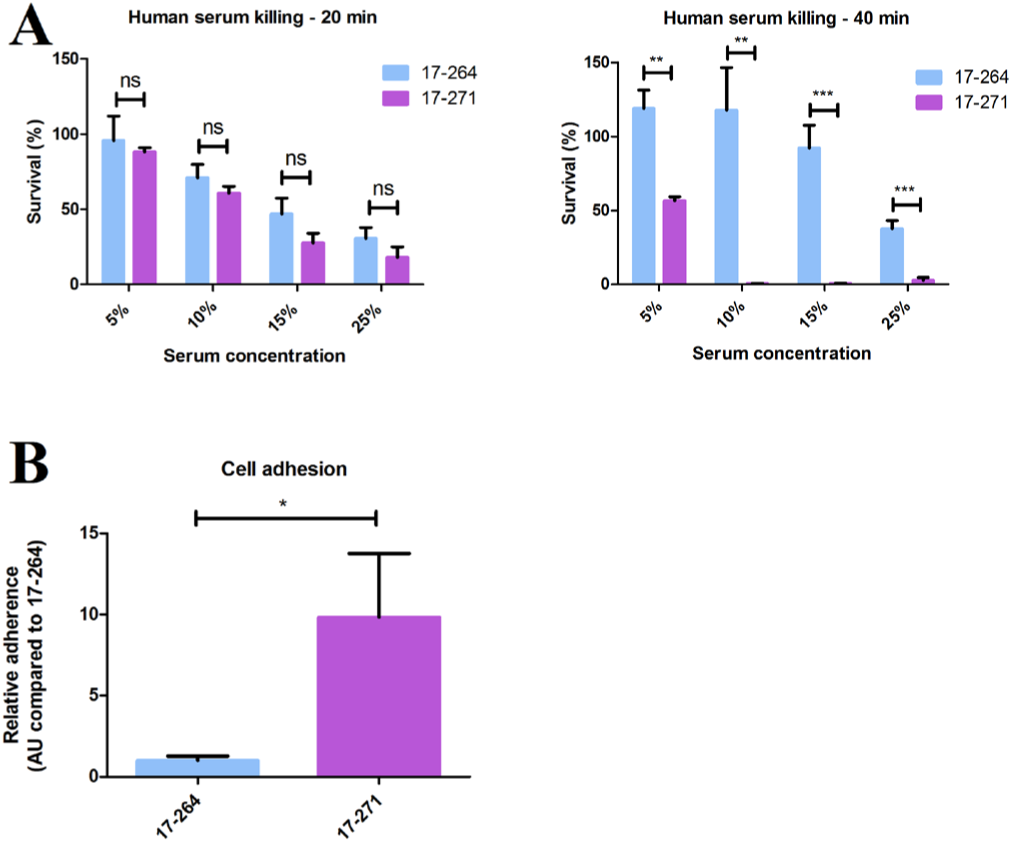
Human serum killing- and cell adhesion assays. A) Human serum killing assay at various concentrations of serum (5-25%) with 20 minutes of exposure (left) and 40 minutes of exposure (right). After 20 minutes, the invasive isolate 17-264 showed a higher survival trend compared to the carrier isolate 17-271. After 40 minutes exposure, 17-264 was able to grow in the various serum concentrations whereas 17-271 could only sustain the 5% serum. Statistical significance values are 5%; *p*=0.0013, 10%; *p*=0.0037, 15%; *p*=0.0004, 25%; *p*=0.0008. B) Cell adhesion assay using the human pharyngeal cell line Detroit-562. The graph was normalised to the adherence of the invasive isolate 17-264. The carrier isolate 17-271 is approximately 10 times better in adherence compared with the invasive isolate 17-264, *p*=0.045. Data information: All graphs are presented with mean ± SEM. *P≤0.05 (Student’s t-test). Graphs and statistical analysis were generated using GraphPad Prism.

These findings from the Swedish isolates prompted the question on growth temperature in the invasive isolate 17-264, while the carrier isolate 17-271 retain a temperature dependent production of capsule (Fig 1E – top). A fraction of the polysaccharide that are the constituents of the capsule are known to be released or “shed” into the surrounding environment by meningococci, when not recycled by the bacterium (15). The increased capsule expression in 17-264 is also evident when observing greater amount of shed capsule from liquid growth at 37°C compared to the 17-271 (Fig 1E – bottom).

To test if the hypercapsulation will improve bacterial survival when facing human immune factors, the two meningococcal isolates were subjected to serum killing assay using pooled serum from healthy individuals. Exposing bacteria to different serum concentrations revealed a trend of higher survival for the invasive isolate 17-264 compared to the carrier 17-271 after 20 minutes (Fig 2A - left). The difference becomes evident after 40 minutes of serum exposure, where the carrier isolate 17-271 was not able to resist 40 minutes serum exposure effectively in any higher serum concentration than 5%, while the invasive isolate 17-264 is able to survive in all serum concentrations, as well as growing more than the input colony forming unit (cfu) (Fig 2A - right). The hypercapsulation phenotype allows the bacteria to resist complement serum killing, however capsulation could inadvertently affect the ability of the bacteria to interact and adhere to pharyngeal cells due to inaccessible surface protein masked by the capsule (16). To investigate this, adhesion assays on human pharyngeal whether the hypercapsulation phenotype due to the loss of an 8bp repeat in the 5’-UTR-*cssA* is a common phenomenon and linked to cases of invasive infection. To address this, the 5’-UTR-*cssA* of meningococcal isolates reported as “invasive” or “carrier” in the PubMLST database from Europe during 1^st^ of January 2010 to 31^st^ of December 2018 was compiled and investigated. The geographical designation of Europe and time span were chosen due to the overall improved WGS and extensive cases reported in the PubMLST database, allowing a sufficient, large, and relevant dataset to be used for analysis. In addition, the two isolates in this study were associated with two European countries; Sweden and France. A total of 4325 invasive- and 176 carrier isolates fitting the search criteria were analysed. Sequences of their 5’-UTR-*cssA* were divided into the three known configurations; two times 8bp repeat (TATACTTATATACTTA), one 8bp (TATACTTA) and one 8bp with substitutionary mutations of either (TAT<u>G</u>CTTA/TAT<u>G</u>C<u>C</u>TA) that would revert capsule production back to the native two times 8bp repeat level. Our statistical analysis shows that the presence of the two times 8bp repeat is similar for both groups with 75% (n=176) in carrier isolates and 73.2% (n=4325) for invasive isolates. Interestingly, the average occurrence of the single 8bp repeat (hypercapsulation) is only 5.1% in the carriers whereas the frequency is more than threefold higher at 16·3% in invasive isolates (Fig 3A). Inversely, the sequences with compensatory mutation revert to almost two fold higher frequency in carrier isolates at 19.9 % compared to the invasive isolates at 10.5%. χ^2^- test was performed and the observed difference in frequency is statistically significant (*p*=0.015).

**Fig. 3.**
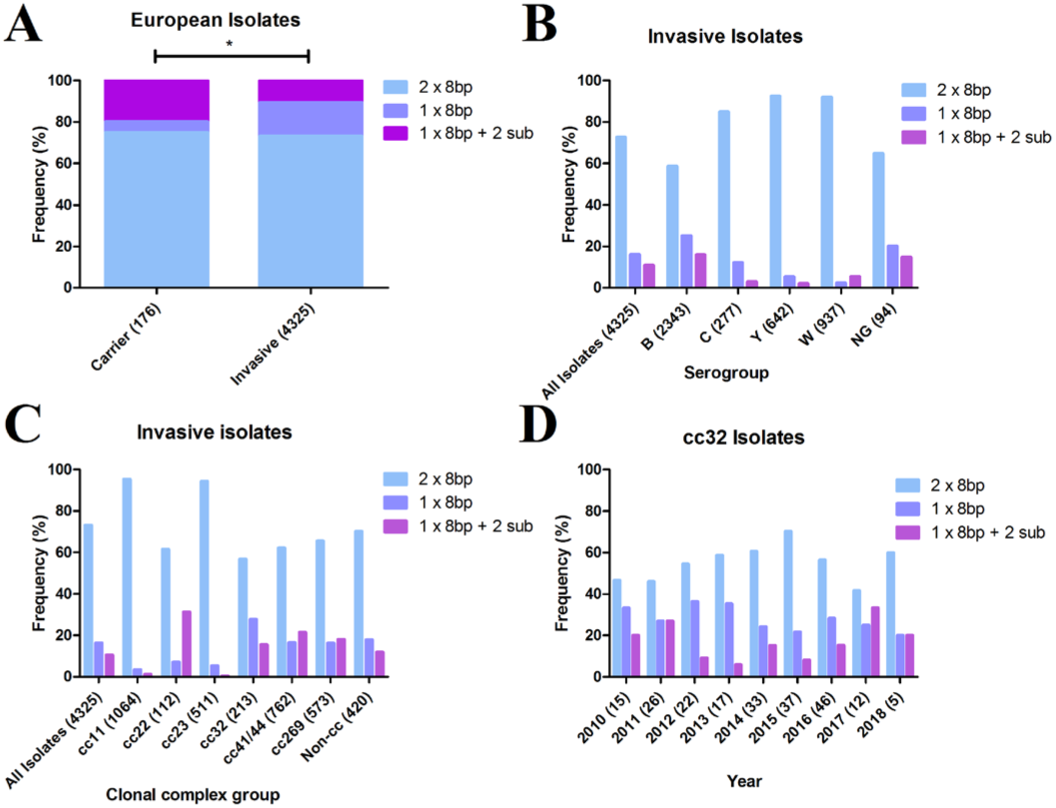
Frequency distribution of 8 bp tandem repeats among 4501 meningococcal isolates in Europe from 2010-2018. A) Frequency distribution showing a three times higher frequency of the 1 x 8bp tandem repeat in the invasive isolates (16.3%) compared to carrier isolates (5.1%). The sequences with compensatory mutation revert to higher frequency for carrier isolates (19.9 %), and lower for invasive isolates (10.5%). χ^2^-test shows significant difference between the distributions (significance value of p=0.015). Post-hoc testing with Bonferroni correction revealed both the 1×8bp (p=0.00007) and 1×8bp+2sub (p=0.0002) distributions to be significantly different between the carrier and invasive isolates. B) The distribution of 8bp tandem repeats among invasive isolates grouped into serogroups display the highest frequency of 1 x 8bp tandem repeat in serogroup B (25.2%) compared to other serogroups (2.5 – 12.3%). C) Distribution of tandem repeats in different clonal complexes of invasive isolates shows that cc32 has the highest proportion (27.7%) of 1 x 8bp tandem repeat. D) Annual distribution of tandem repeats of cc32 reveals consistent frequencies throughout the years of 1 x 8bp repeat (range 20.0– 36.4%). Data information: Amount of isolates analysed are designated by n. *P≤0.05 (χ2-test), *P≤0.0083 (Bonferroni corrected). All graphs and statistical analysis were generated using GraphPad Prism and SPSS. Numbers in brackets denotes number of isolates analysed.

The majority of the European invasive isolates investigated in this study are of serogroup B, with higher distribution of a single 8bp tandem repeat (25.2%, n=2343) than the other serogroups respectively, C (12.3%, n=277), Y (5.5%, n=642), W (2.5%, n=937) and in non groupable (NG) (20.2%, n=94) (Fig 3B). As specific ccs are more prevalent in invasive disease, the configurations of 5’-UTR-*cssA* of invasive isolates were also divided into respective complexes (Fig 3C). The same cc (cc32) as the investigated clinical isolates used in this study showed an even higher frequency with the loss of an 8bp repeat, 27.8% (n=213), than the overall European invasive isolates of 16.3%. To investigate whether the high frequency of the loss of an 8bp repeat among cc32 isolates is a recent occurrence, the cc32 data were also subdivided into years of incidence (2010-2018). The loss of an 8bp repeat frequency remains high throughout the years (Fig 3D).

## Discussion

Our work supports a regulatory RNA to be directly involved in the clinical manifestation of meningococcal disease. Epidemiological analysis of 4501 meningococcal isolates reveal that the loss of an 8bp tandem repeat in this regulatory RNA is more than three times frequent in invasive clinical isolates thus emphasising its role in virulence. Molecular analyses from the two Swedish isolates in this study have further confirmed our previous finding that the 8bp tandem repeats in the 5’-UTR-*cssA* regulate CssA expression in a temperature dependent manner (11). In addition, we demonstrated that the higher expression of CssA leads to higher production of capsule as shown by the dot blot assays.

It is known from previous studies that both expression levels of T4P and capsule affects the ability of meningococci to adhere to human cells (17, 18). Our results indicate that while the invasive isolate 17-264 express more PilE protein, the most abundant minor pilin in the makeup of the T4P it still has a reduced ability to adhere to cells. Our findings supports previous research that the thicker polysaccharide capsule could interfere with important surface structures such as T4P thus sequestering their function for adhesion to pharyngeal cells (19). Previous study has shown that T4P and Opa mediates endothelial interactions in a synergistic manner in capsulated strains (20). It is also known that expression of PilE and Opa is in unison to facilitate Neisserial microcolony formation (21).

Our analysis here also show nine copies of incomplete *IS1655* elements in the carrier isolate, 17-271. All of the *IS1655* hits were sequenced at the end of contigs in the WGS data and with a short alignment to the reported *IS1655*. This *IS* element is an exclusive feature of meningococcus and found in high numbers in several strains (22, 23).

To date, no work has been done to investigate whether *IS1655* is involved in meningococcal pathogenesis, given its prominent and diverse presence across the meningococcal genomes.

In our previous study investigating this 8bp tandem repeats (11), we speculated that the three configurations of the 8bp tandem repeats are an evolutionary step-by-step process. The native two 8bp tandem repeats and temperature regulated capsule expression are beneficial for the bacterium during non-invasive colonisation of the host nasopharynx. Meningococci face environment switches constantly, such as localised inflammation caused by the bacterium itself, other microbes, or by entering the blood circulation. It is our hypothesis that the stress induced by high amounts of immune factors together with a febrile condition will make the thermostable two 8bp repeats superfluous, as the need for protection by expressing more polysaccharide capsules is great. This selection pressure induced by the host alters the meningococcal population through the loss of an 8bp repeat in the 5’-UTR*-cssA* to produce more capsule constitutively. The rapid onset of the invasive disease together with the presence of all three different 5’-UTR-*cssA* thermosensors in carrier isolates suggest this selection to take place in the nasopharynx. To investigate the loss of 8bp mechanism, we subjected the carrier isolate 17-271 to a sub-lethal concentration of 6% human serum stress. After two and a half hours, we were able to identify colonies that have lost a single 8bp (Fig S3). Stress induced modification of tandem repeats have previously been shown to regulate other immune evasion factors such as the meningococcal PorA outer membrane protein (24) and the gonococcal iron-repressible protein FetA (25). These tandem repeats are prone to strand-slipping during DNA replication especially during stress condition as a form of bacterial selection strategy to the new environment (26). During recovery conditions after stress, the meningococci would shift towards a population that restores colonisation by reducing capsule expression with a reinstated thermostable 5’-UTR-*cssA* by one or two point mutations to strengthen the stem-loop structure (Fig S4). Interestingly, among the 4501 meningococcal isolates, we could observe this phenomenon as an almost two-fold higher frequency of one 8bp tandem repeat with substitutionary mutations among the carrier isolates compared to the invasive isolates. Capsular switching is a phenomenon where meningococci acquire parts of the *cps* locus from other serogroups such as B to C or the reverse (27). The serogroup C, cc-32 isolates used in this study could support a capsule switching event prior to the alteration of the 5’-UTR-*cssA* as similar capsular switching has been reported previously (28, 29). It is interesting to hypothesise how capsular switching could also contribute to the “reversal” of a 1×8bp or 1×8bp + 2sub 5’-UTR-*cssA* to a “native” 2×8bp 5’-UTR-*cssA* as a consequence of capsular switching bringing the 5’-UTR-*cssA* of the “donor” DNA, however previous work has shown the intergenic region between *ctrA* and *cssA* is not included in capsular switching (27).

While it is difficult to determine all the host genetic- and environment factors involved, we postulate that the loss of an 8bp tandem repeat hypercapsulation phenotype of the 17-264 isolate contributed to the fatal disease manifestation in the adolescent girl from Sweden. Statistical analysis further strengthen this hypothesis as the one 8bp tandem repeat is three-fold more frequent among invasive isolates compared to the carrier isolates. Our epidemiology data also show the prevalence of the loss of an 8bp tandem repeat is most prominent in Serogroup B, cc32. In addition we demonstrated that invasive isolates of other cc such as cc41/44 regulate CssA expression solely depending on the configuration of the 8bp tandem repeats (Fig S5) regardless of cc. It is important to note that the isolation of clinical isolates are from two distinct sites; the nasopharynx (carrier) and blood/cerebrospinal fluid (CSF) (invasive). The presence of all three configurations of the 8bp tandem repeats in both sites reflect the importance of this regulatory RNA in controlling its capsular biosynthesis in both environments. However, the significant higher prevalence of the hypercapsulated single 8bp configuration observed among the invasive isolates found in blood/CSF would be beneficial for the bacteria to evade the host immune killing. It is hard not speculate that while the single 8bp background could be due to isolation sites, our statistics finding emphasize that this regulatory RNA plays a pivotal role in virulence and is associated meningococcal disease manifestation.

Due to the lack of suitable infection models, we believe that molecular comparative WGS between meningococcal isolates from invasive and carrier cases is the best alternative. With WGS, polymorphism comparisons should include regions such as 5’-UTRs and regulatory RNAs as they are significantly involved in the expression of many virulence factors exemplified here by capsule production. Comparative WGS will facilitate future studies into underlying mechanisms involved in disease manifestation, improve diagnostic techniques and treatments by clinicians and advance future vaccine development.

## Materialsand Methods

### Bacterial isolates and DNA extraction

The bacteria are cultured overnight on chocolate agar in 37°C and 5% CO_2_, serogrouped by coagglutination, and stored at −70°C. In this study, both meningococcal isolates were isolated during May 2017 and whole-genome sequencing (WGS) was performed. DNA was extracted using a customized protocol and QIAsymphony DSP virus/pathogen Midi kit on the QIAsymphony system (Qiagen, Hilden, Germany). WGS was performed using the Nextera XT DNA library preparation kit, the MiSeq reagent kit v3, 600 cycles on the Illumina MiSeq platform (Illumina, San Diego, CA). The reads were assembled *de novo* using Velvet (30). The quality of sequencing was controlled by the N50 value, contig count and coverage. The sequences were trimmed until the average base quality (Phred score) was >30 in a window of 20 bases. The assemblies were uploaded to the *Neisseria* PubMLST database, and the sequences were automatically scanned and tagged against defined “NEIS” loci in the database (31). Information regarding the typing of the French isolate was obtained through personal communication with Dr. Muhamed-Kheir Taha, Institut Pasteur, Paris

### Bacterial culturing

For all other purposes, bacteria were grown with Brain Heart Infusion (BHI) broth (Sigma-Aldrich, Saint-Louis, MO) or agar supplemented with horse serum (Thermo Fisher Scientific, Waltham, MA). For all experiments, bacteria were inoculated on BHI agar for 16h, in 37°C and 5% CO_2_. Liquid cultures were grown in BHI broth starting at an optical density at 600 nm (OD_600_) of 0·05 and cultured to OD_600_:0·6 where protein or DNA were harvested.

### Scanning Electron Microscopy

Bacteria were fixed with 6% paraformaldehyde and transferred to a pre-sputtered filter (Polyamide, NL 16, GE Healthcare UK Limited, Buckinghamshire, UK), rinsed in distilled water and placed in 70% ethanol for 10 minutes, 95% ethanol for 10 minutes and absolute ethanol for 15 minutes, all at 4°C and then into acetone. Specimens were then dried using a critical point dryer (Balzer, CPD 010, Lichtenstein) with CO_2_. After drying, filter was mounted on an aluminium stub and coated with Platinum (Q150T ES, West Sussex, UK). The specimens were analysed in an Ultra 55 field emission scanning electron microscope (Zeiss, Oberkochen, Germany) at 5 kV.

### Immunoblotting and dot blotting

For immunoblotting, bacterial pellet was spun down at 7,000 g for 10 minutes and lysed in Buffer A (200mM KCl, 50mM Tris-HCl pH 8·0, 1mM EDTA, 10% Glycerol). Cell debris was spun down at 12,000 g for 5 minutes and lysate collected for protein quantification. using BCA (Thermo Fisher Scientific). SDS-loading buffer was added and boiled at 98°C for 10 minutes before protein was subjected to electrophoresis on 4-12% Bis-Tris gel (Thermo Fisher Sceintific) and run at 110V. Gels were transferred to PVDF membranes using the Trans-Blot Turbo system (Bio-Rad, Hercules, CA). For dot blots, bacteria were harvested from plates grown at 30-, 37-, and 42°C, and supernatant was harvested from liquid growth at 37°C. Supernatant was sterile filtered and serial diluted in PBS while harvested bacteria were lysed as described previously and diluted 1×10^-4^ in PBS. Amersham Hybond-P PVDF membranes (GE healthcare, Chicago, IL) were activated and 10µl of the samples were dotted onto the membrane and left to dry for one hour at 37°C. Membranes were blocked in 5% milk for one hour at room temperature (RT) and incubated with primary antibody/anti-sera in PBS + 0·1% Tween, for one hour at RT. After washing membrane four times 5 minutes, secondary antibody was added in PBS + 0·1% Tween for one hour RT. Membranes were exposed by Amersham ECL reagents (GE healthcare) and visualised by GelDoc XRS+ (Bio-Rad). Commercial antibodies used was anti-RecA, (Abcam, ab63797) (crossreacts with *N. meningitidis* RecA), anti-PorA P1.7, (NIBSC, 01/514) and anti-fHbp, (NIBSC, 13/216). The remaining antibodies used are custom antibodies obtained from Yvonne Pannekoek (anti-Hfq), Hank Seifert (anti-PilQ), Ann-Beth Jonsson (anti-PilC) and Ryoma Nakao (anti-Rmpm, -PilE, -Opa, -CssA and GroEL).

### Human serum stress and killing assay

Human serum killing assay was performed with bacteria resuspended in PBS and diluted to a concentration of 1×10^5^ CFU/ml in DMEM-medium (Gibco, Thermo Fisher Scientific). Serum from healthy humans (Sigma-Aldrich) was used to stress bacteria at 37°C in the presence of 5% CO2 for 20-, or 40 minutes (killing assay) or hours (serum stress induction). Survival of bacteria in the presence of sera was determined by dot-plating 10µl and counting CFUs after overnight incubation. The percentage of survival was measured relative to input.

### Bacterial adhesion assay

Detroit 562 cells were cultured in DMEM, supplemented with 10% FBS (GE Healthcare). 5×10^5^ cells were seeded per bacterial isolate tested. Cells were washed with PBS and covered with 2mL DMEM without FBS. Bacteria were resuspended in PBS to an OD_600_:0.5 and added to the cells at a MOI of 50. Bacteria and cells were incubated at 37°C at 5% CO_2_ for one hour, and washed three times with PBS. Cells were lysed for 10 min with 1% saponin (Sigma-Aldrich), and the lysate was diluted 1:3 in PBS before dribble-plating and incubated at 37°C, ON and analysed the next day. CFU of the negative control (no cells) were subtracted from CFU of the output. The result was normalised to the CFU of the input.

### Whole-genome sequencing with loci comparison

17-264 and 17-271 isolates were compared using the “Genome comparator” function in the PubMLST database. Sequence alignments were performed for all loci that were marked as different between the isolates. For further protein sequence prediction and alignment, online tools; ExPASy and Clustal Omega (SIB Swiss Institute of Bioinformatics, Lausanne, Switzerland) were used.

### 5’-UTR-cssA mapping and in silico RNA structure predictions

The PubMLST database was used for identifying meningococcal isolates and selected according to the following search criteria. The range from the 1^st^ of January 2010 to 31^st^ of December 2018 and geographic restriction to the continent of Europe. The searches were then separated by the designation of “carrier” or “invasive (unspecified/other)”. Hits were BLAST to a 200bp long sequence that encompasses the 5’UTR of the *cssA* gene and the beginning of the coding region of *cssA*. The results were exported and analysed by excluding all hits that had less than 100bp alignment to the search query. Any isolate that was suspect of non-canonical 5’UTR-*cssA* or where the sequence could not be retrieved, was excluded from the analysis. To predict, analyse, and visualise secondary structure of RNA, multiple available softwares were used. RNAfold by ViennaRNA package (32) and mFold (33) were used for structure prediction data and folding based on the minimum free energy. VARNA (34) was used to visualise the secondary structure obtained together with annotated nucleotides.

### Statistical methods

All experiments were performed with two or three biological replicates. Data from human serum killing assay and bacterial adhesion assay are shown with mean ± SEM, and students t-test was used to calculate statistical significance. Normal distribution are assumed for this data. The observational data of the 5’UTR-*cssA* mapping is shown with mean distribution and χ^2^-test was performed to calculate statistical significance in group distributions. P values of less than 0.05 was considered statistically significant. Graphpad Prism v5.04 (Graphpad software, Inc, La Jolla, CA) and SPSS v25 (IBM, Armonk, NY) was used to plot data and perform statistical analysis.

## End Matter

### Author Contributions and Notes

S.J. performed the meningococcal isolates isolation, typing and WGS. J.K., H.E. and E.L. performed the experiments and analysed the data. J.K. performed the European clinical data collection and analysis. E.L. provided overall direction. E.L and J.K. performed literature search and wrote the manuscript with input from H.E. and S.J.

The authors declare no conflict of interest.

This article contains supporting information online.

## Acknowledgments

We thank Dr Francesco Righetti and Dr Kristiina Tammimies for critical reading of the manuscript. Most of the non-commercially available antibodies used were gifts from Dr Ryoma Nakao from The National Institute of Infectious Diseases at Tokyo, Japan. This publication made use of the *Neisseria* Multi Locus Sequence Typing website (https://pubmlst.org/neisseria/) developed by Keith Jolley and sited at the University of Oxford (35). The development of this site has been funded by the Wellcome Trust and European Union. This study also made use of the Meningitis Research Foundation Meningococcus Genome Library (http://www.meningitis.org/research/genome) developed by Public Health England, the Wellcome Trust Sanger Institute and the University of Oxford as a collaboration, a project funded by Meningitis Research Foundation. This work was funded by the Swedish Foundation for Strategic Research (ICA14-0013), Knut and Alice Wallenberg Foundation, and Swedish Research Council (Dnr: 2014-2050).

