## Supplemental Figures for "A regulatory RNA is associated to invasive meningococcal disease in Europe"

**Supplementary Figures**

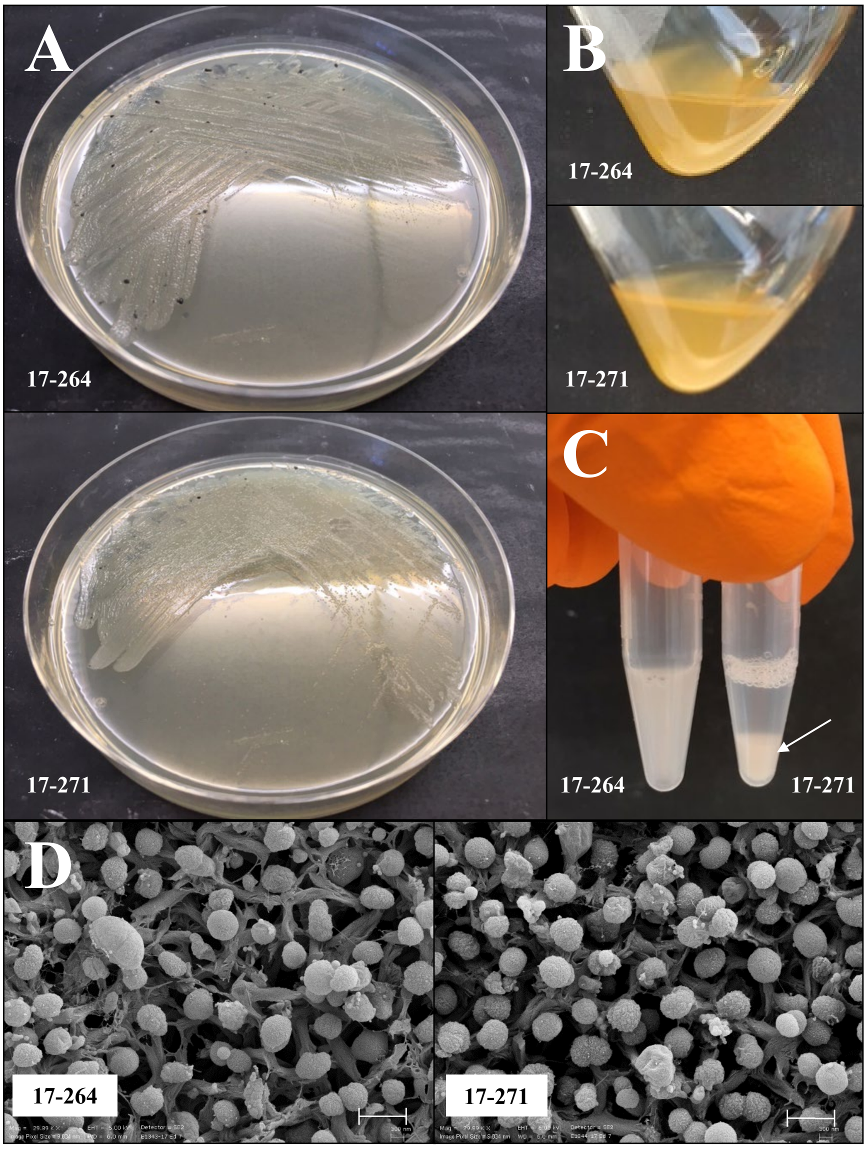


**Figure S1**.

A) Overnight growth on BHI agar shows no gross difference in colony morphology.

B) Growth in liquid BHI broth with shaking, 180 rpm, shows no apparent differences.

C) Static suspension of bacteria in PBS reveal sedimentation of the carrier isolate 17-271 after 15 minutes (arrow).

D) Scanning Electron Microscopy (SEM) of the two different isolates shows no distinct morphological difference. Scale bar denotes 1 µm.

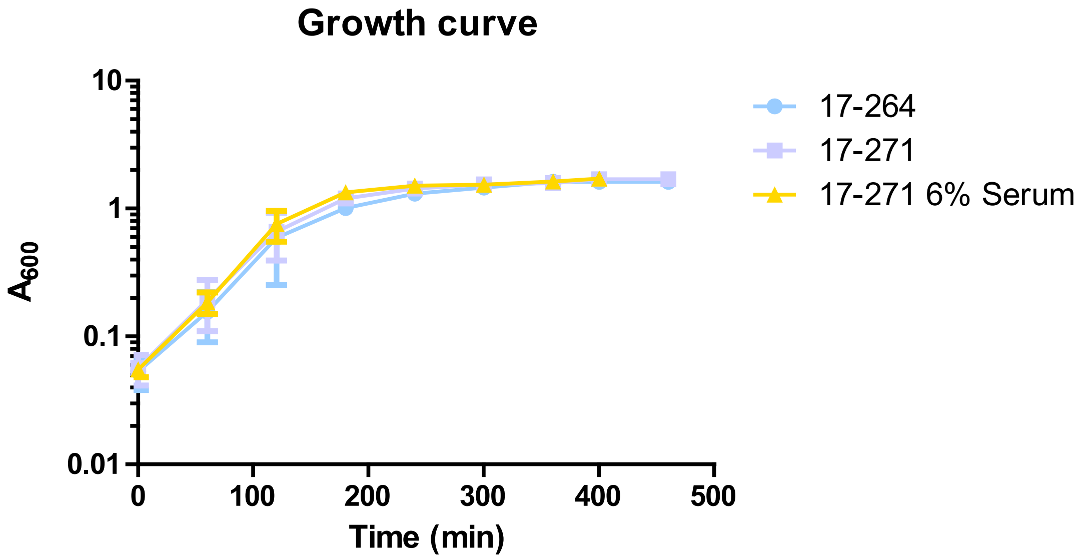


**Figure S2.**Growth curves of invasive isolate 17-264, carrier isolate 17-271, and 6% serum-stressed isolate17-271 show similar growth patterns.

**A**

```
                 +1
17-264           GGCCTAATTATAGCACTTAATCGAAATAAATTTATGAGTACGTAGAGTATAATTAGTATTCTTCTTTCCAACTTCCTTATACTTA--------TAGATTCTAAAATCATG
17-271           GGCCTAATTATAGCACTTAATCGAAATAAATTTATGAGTACGTAGAGTATAATTAGTATTCTTCTTTCCAACTTCCTTATACTTATATACTTATAGATTCTAAAATCATG
17-271 6% Serum   GGCCTAATTATAGCACTTAATCGAAATAAATTTATGAGTACGTAGAGTATAATTAGTATTCTTCTTTCCAACTTCCTTATACTTA--------TAGATTCTAAAATCATG
```

**B**

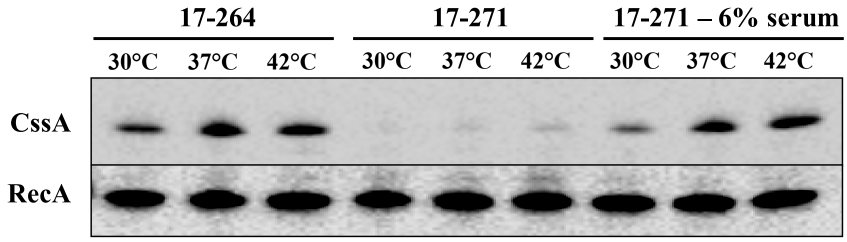


**Figure S3.**
A) Sequence alignment after 6% serum exposure reveals the loss of an 8 base-pair tandem repeat in the 5´-UTR-*cssA* of carrier isolate 17-271. The tandem repeat, RBS and start codon are marked in red/green, cyan and yellow, respectively. Small blue dots designate base pairing.

B) Immunoblot showing 6% serum induced the loss of an 8bp in the previous 2 x 8bp tandem repeat 17-271 isolate, resulting in high CssA expression.

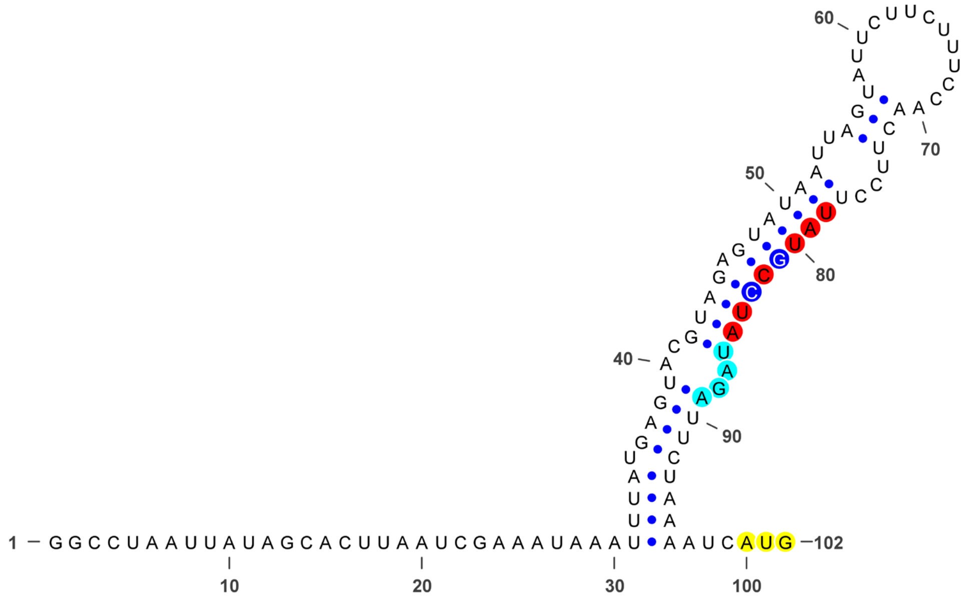


**Figure S4.** Secondary structure prediction of the 1 x 8bp + 2 substitutions showing a restored stem-loop structure.

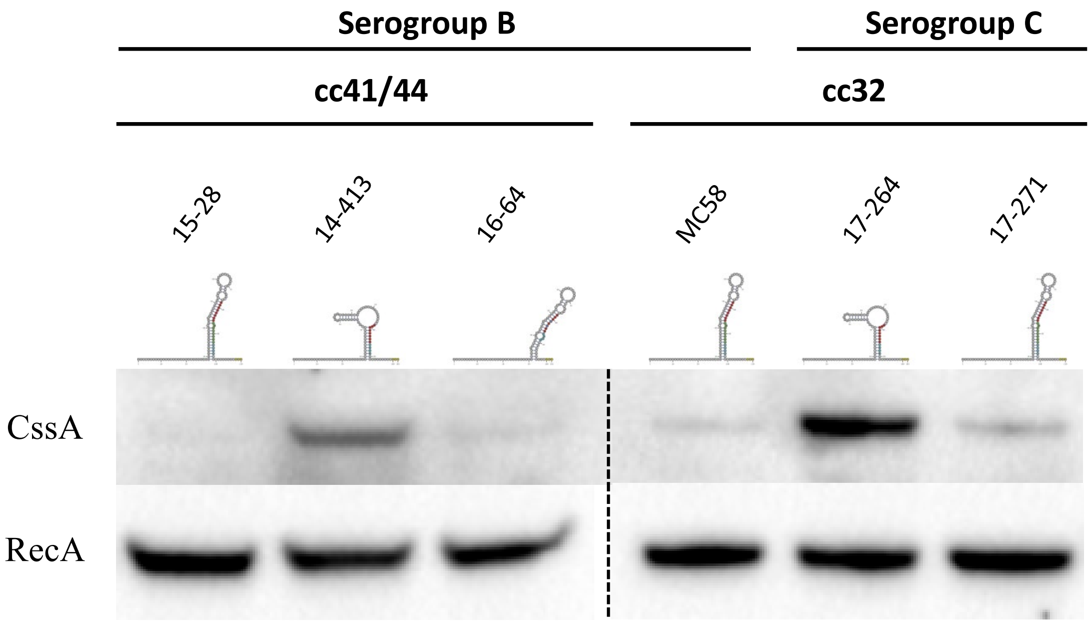


**Figure S5.** Immunoblot of CssA expression of different meningococcal isolates, including the serogroup B, *N. meningitidis* MC58 as well as representative clinical isolates with the three configurations of the 5´-UTR-*cssA*.
